# A high-throughput approach for the efficient prediction of perceived similarity of natural objects

**DOI:** 10.1101/2024.06.28.601184

**Authors:** Philipp Kaniuth, Florian P. Mahner, Jonas Perkuhn, Martin N. Hebart

**Author notes:** Correspondence should be addressed to Philipp Kaniuth or Martin Hebart.

## Abstract

Perceived similarity offers a window into the mental representations underlying our ability to make sense of our visual world, yet, the collection of similarity judgments quickly becomes infeasible for larger datasets, limiting their generality. To address this challenge, here we introduce a computational approach that predicts perceived similarity from neural network activations through a set of 49 interpretable dimensions learned on 1.46 million triplet odd-one-out judgments. The approach allowed us to predict separate, independently-sampled similarity scores with an accuracy of up to 0.898. Combining this approach with human ratings of the same dimensions led only to small improvements, indicating that the neural network used similar information as humans in this task. Predicting the similarity of highly homogeneous image classes revealed that performance critically depends on the granularity of the training data. Our approach allowed us to improve the brain-behavior correspondence in a large-scale neuroimaging dataset and visualize candidate image features humans use for making similarity judgments, thus highlighting which image parts may carry behaviorally-relevant information. Together, our results demonstrate that current neural networks carry information sufficient for capturing broadly-sampled similarity scores, offering a pathway towards the automated collection of similarity scores for natural images.

## INTRODUCTION

Understanding the nature of human mental representations is a key aim of the cognitive sciences^1^. A central approach for gaining access to these representations has been the collection of perceived similarity judgments^2^, with a long history across the fields of psychology, linguistics, neuroscience, and computer science^3–6^. Similarity judgments allow us to improve our understanding of a variety of cognitive processes, including object recognition, categorization, decision making, and semantic memory^6–13^. In addition, they offer a convenient means for relating mental representations to representations in the human brain^14,15^ and other domains^16,17^.

To collect similarity judgments, a diverse set of behavioral similarity tasks has been developed, including pairwise ratings^18–20^, pile sorting^21,22^, object arrangements^23–25^, triplet judgments^24,26–28^, or speeded, more indirect similarity tasks^29–31^. While each task has its unique advantage, a key challenge with the acquisition of similarity ratings is their complexity, which in many cases grows quadratically or cubically with the number of items (superlinear complexity, Figure 1). This challenge not only limits our understanding of the similarity between images to small stimulus sets; it also limits the direct comparability between studies that rely on different images. A more efficient approach for sampling similarity scores mirroring these judgments is thus highly desirable and could even be applied retrospectively to a large number of existing datasets that did not collect similarity scores. By overcoming the data sampling challenge, such an approach would thus open novel data-driven and hypothesis-driven avenues for understanding mental representations at scale.

**Figure 1.**
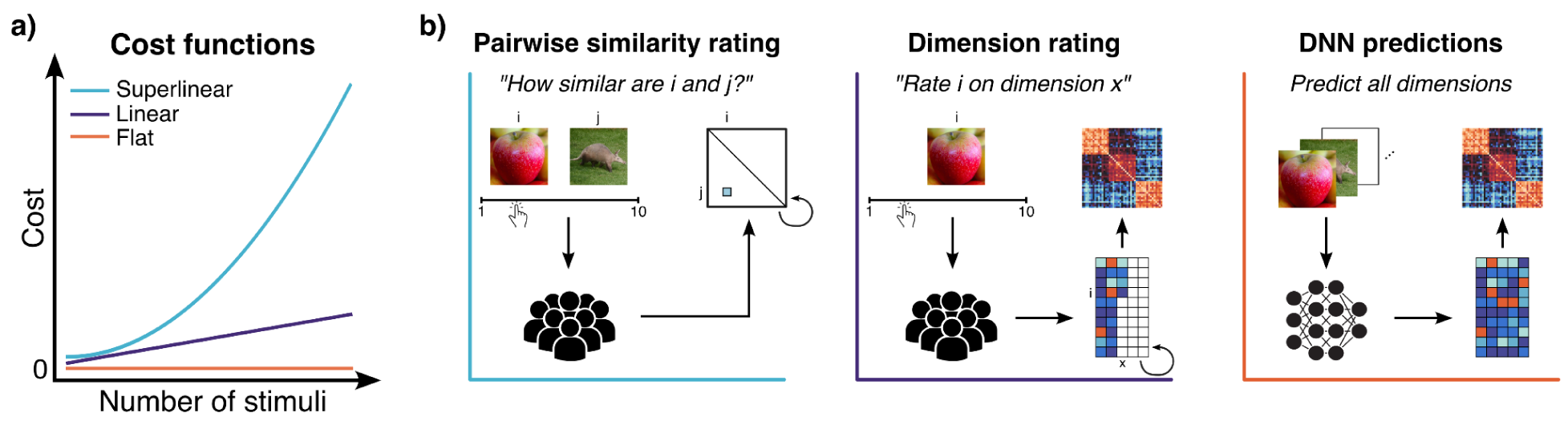
Cost functions and different perceived similarity sampling regimes. (a) For a stimulus set with *n* stimuli (x-axis) the cost of sampling all similarities (y-axis) grows differently for different sampling regimes. (b) For most classical behavioral similarity tasks like the pairwise rating task (left), cost grows superlinearly. Ideally, though, behavioral tasks should have linear cost increases like the task we present in this study (middle). Flat costs (right) are only attainable when utilizing deep neural networks (DNNs) and applying an automatic method like the one we explore in this study with which the whole embedding for a given image set is predicted in one go which can then be used to compute the whole representational similarity matrix.

An efficient alternative to directly sampling similarity judgements is offered by feature-based rating approaches (Figure 1b, middle). Here, stimuli are first scored along multiple dimensions that can be both visual (e.g., shape features) or semantic (e.g., animacy) in nature. Then, those dimensions are combined to compute an overall similarity score between pairs of stimuli^13,32–34^. While this leads to increased efficiency that scales linearly with the number of items (Figure 1a), the predictive accuracy of this approach has been rather limited^33,35, but see 36^. More recently, computational models have been used (Figure 1b, right), based on which predictions of similarity can be automated, including the use of raw features of computational models^35,37^ or the indirect use of large language models^38–40^. Similar to behavioral feature ratings, prediction accuracy of these approaches has remained limited or was restricted to small-scale comparisons, and comparisons were often done on image sets where the tested categories in the training and testing set overlapped^35,38,39,41^. This leaves open the question to what degree results would generalize to a broader set of independent stimuli^35,39,42,43^.

To overcome existing challenges of predicting perceived similarity judgments, here we present an efficient automated approach that is based on predicting a small set of human interpretable dimensions derived from similarity ratings to 1,854 diverse object images. This approach allows us to retrieve similarity scores, which we can validate with independently sampled behavioral ground-truth similarity. We showcase our approach on different image sets across various neural network models. Our results show that the approach is capable of generating accurate perceived similarity scores for various broadly-sampled stimulus sets. Combining this automated approach with human judgments of the same dimensions led only to small improvements, indicating that the neural network relied on largely similar information as humans for predicting representational dimensions. Having established an end-to-end mapping of input images to perceived similarity that generalizes to a large set of natural images, we showcase the potential of this approach by (1) revealing image regions that are predicted to be most relevant for similarity judgements and (2) improving the brain-behavior correspondence in a large-scale neuroimaging dataset. Together, the results demonstrate that the prediction of interpretable representational dimensions offers a powerful approach for the efficient prediction of perceived similarity, opening the avenue to automatically sample perceived similarity scores for broadly-sampled image categories.

## RESULTS

To automate the generation of similarity judgments, we tested the degree to which off-the-shelf deep neural networks (DNNs) contain representations sufficient for capturing perceived similarity. Instead of directly using raw DNN features to predict similarity^41^, we developed a two-step procedure (Figure 2, top). In the first step, we predicted a human interpretable similarity embedding for each stimulus from DNN representations of these images. In the second step, we derived perceived similarity scores from this predicted embedding. For the first step, as a similarity embedding we used the Sparse Positive Similarity Embedding (SPoSE), which consists of 49 human interpretable dimensions^36^. SPoSE had been trained on 1.46 million behavioral triplet odd-one-out judgments across 1,854 object images from the THINGS database, a sizable database containing thousands of images that comprehensively cover objects that are nameable in the American English language^44^. The behavioral dataset has been shown to accurately predict similarity judgments in held-out data close to the noise ceiling^36^, making it ideally suited for the prediction of perceived similarity. To predict the similarity embedding, we extracted DNN activations for each of the 1,854 reference images. Next, we trained a linear encoding model on this DNN embedding to predict values on each of the 49 dimensions separately. By breaking down the problem of predicting stimulus similarity into smaller units, we allow the DNN to focus on one dimension at a time rather than having to account for all possible dimensions in parallel. Pairing this approach with a large-scale similarity dataset offers a promising avenue for predicting embedding dimensions with high precision.

**Figure 2.**
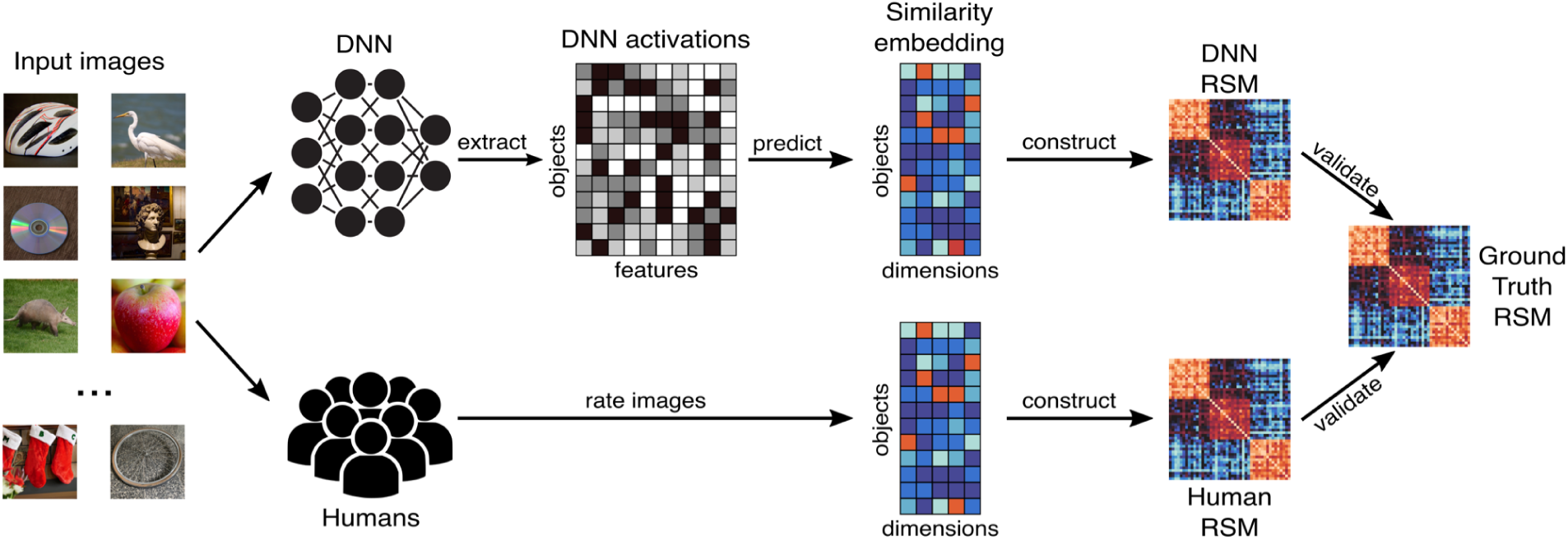
Overview of our approach. For a given image set, we extracted image activations from various image-computable neural network models (top). Based on these activations, we trained an L2-penalized multi-output multiple regression encoding model to predict a 49-dimensional Sparse Positive Similarity Embedding (SPoSE). We then used this embedding to construct a DNN-predicted representational similarity matrix (RSM) which we validated with a behavioral ground-truth RSM. To compare our DNN-based approach to human performance, we conducted crowdsourcing to receive human-generated ratings of all images on the same embedding (bottom). Based on this embedding, we computed a human-constructed RSM which we validated with the ground-truth RSM. Images in this figure were taken from the THINGSplus dataset and are used for illustrative purposes only^79^.

Having predicted the similarity embedding from DNN activations, in the second step we used these predicted dimensions to construct perceived similarity between all pairs of stimuli. To achieve this, we mimicked the cognitive process of constructing similarity judgments assumed for the generation of SPoSE similarities. For a given pair *i* and *j*, similarity was constructed by computing all possible triplet odd-one-out judgments from the embedding involving both stimuli and all possible contexts imposed by a third stimulus *k*. Across all possible *k*, similarity for *i* and *j* was defined as the probability of predicting stimuli *i* and *j* as belonging together and stimulus *k* being the odd-one-out. Repeating this procedure for all possible pairs of stimuli allowed us to construct a predicted representational similarity matrix (RSM), which we could later validate with a ground-truth RSM that we obtained independently.

### Accurate prediction of perceived similarity for broadly-sampled natural object images

To achieve high prediction performance for human similarity, we sampled activations from a broad selection of DNN architectures that had been trained with different objectives and training datasets (see *Methods*). This breadth additionally allowed us to avoid that results were a mere idiosyncrasy of the chosen network and reveal the extent to which the networks vary in their prediction of human similarity judgments. The results of the comparison of DNNs is shown in Figure 3. As a baseline, we used (1) classical representational similarity analysis (cRSA)^17^, which directly computes similarity from DNN embeddings and is an established method for relating DNN representations with perceived similarity, and (2) feature-reweighted RSA (FR-RSA)^16,41,45–47^, which computes similarity from linearly-reweighted features learned from the original similarity matrix and has been shown to achieve much stronger correspondence to similarity judgments^41,45^. Across DNNs, the prediction of perceived similarity varied strongly. However, for a number of DNNs, the predictive performance was excellent, achieving correlations with ground truth of *r* = 0.887 for the visual encoder layer of the multimodal network OpenCLIP-ResNet50 trained on 400 million image-text pairs, and *r* = 0.888 for OpenCLIP-ViT-g-14 trained on 2 billion image-text pairs. All of the best neural networks were based on a variant of CLIP, which indicates the benefit of using large, diverse training data, or a training objective that includes much broader semantics based on text descriptions of images^48–52^. Between the poorer models and the CLIP models was a range of other network architectures with different training objectives, indicating that architecture and objective were not dominant factors at predicting similarity. DimPred clearly outperformed the baseline approaches, yielding improved predictive accuracy for 52 of 53 computational models.

**Figure 3.**
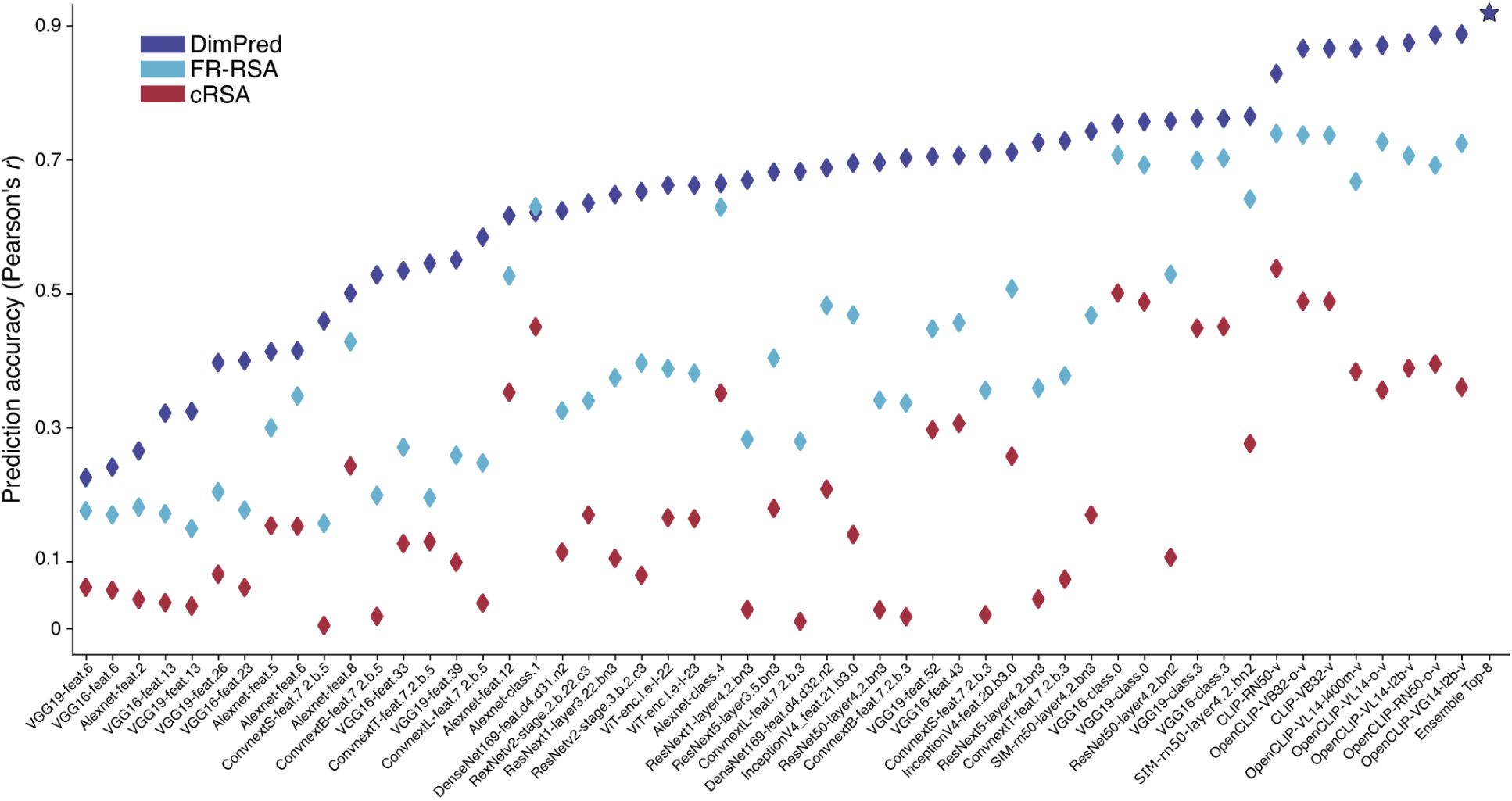
Predictive performance of all computational models for perceived similarity. For the 1,854 reference image set, separately for each computational model (x-axis) we Pearson-correlated the predicted and ground-truth similarity scores (y-axis). Across models, DimPred outperformed established representational similarity methods that relate representational spaces to each other (cRSA and FR-RSA). Amongst all computational models, Open-CLIP models performed exceptionally well. An ensemble DimPred model consisting of the best 8 computational models performed even better (denoted by a star symbol). For a mapping of computational model name abbreviations to full model names please see Supplement 4.

The computational models also successfully predicted individual embedding dimensions (see Supplement 1).

Since many individual models performed well, we wondered to what degree their combination would improve the prediction of similarity. To this end, we combined values predicted for a specific dimension from different computational models and evaluated whether this led to improved predicted similarity scores. Specifically, we combined the predicted values of the best *n* models for each dimension, creating an ensemble model of predicted perceived similarity. The ensemble model exhibited an even greater predictive accuracy than the best model individually, producing correlations of predicted similarity with ground-truth of up to 0.926 when combining the best 8 computational models (for further details, see Supplement 2).

### DimPred generalizes to out-of-set images and other similarity tasks

To test the degree to which the results generalize beyond the distribution of the 1,854 reference images and the exact similarity task used, we sought to test the performance of DimPred by predicting similarity for independent stimulus sets that had been collected with a variety of different behavioral similarity tasks. Predicting similarity for external validation sets is usually not done when applying cRSA or FR-RSA^17,46,53^, which at least in the case of FR-RSA might lead to overestimating generalization performance. In addition, even though we labeled the similarity matrix of the 1,854 images as “ground truth”, it was not sampled empirically but was the result of similarity prediction of the original similarity embedding, which may slightly overestimate the performance of this approach on external data. To address these issues, we applied DimPred to five additional image sets, each consisting of images from various categories. One validation set consisted of 48 images of categories that are part of the 1,854 THINGS categories that DimPred had been trained on, but contained new images, thus testing generalization at the image level (48- image-generalization image set). Another validation set consisted of 48 images, but this time of concepts not included in THINGS, thus testing generalization to new images and new concepts (48-concept-generalization image set). The three other validation sets consisted of various images sampled in previously published data and collected with different similarity tasks, containing 92 images without natural background sampled with the multi-arrangement task (Mur-92)^54–56^, 118 images natural images sampled with the same task (Cichy-118)^15,57^, and 120 images from a mix of categories sampled with a pairwise similarity task (Peterson-various image set)^41^. We again estimated the performance of DimPred across all computational models and included cRSA and FR-RSA as a baseline.

The results are presented in Figure 4, with a direct contrast against the baselines cRSA and FR-RSA. DimPred again showed strong overall predictive performance on both 48 image sets, with predictive accuracy reaching *r* = 0.858, while it was slightly lower in the additional three image sets (with *r* up to 0.809), indicating a slight drop in performance when generalizing across similarity tasks. When applying the ensemble model of predicted perceived similarity with the best training performance for the 1,854 reference image set, predictive accuracy on the 48 image sets reached up to *r* = 0.898, and for the additional three image sets up to *r* = 0.843 (see Supplement 2 for further details). For all image sets, DimPred was clearly superior to the baseline approaches. Together, this demonstrates that DimPred generalizes well to other images as well as other concepts while also generalizing well to other similarity tasks, underscoring the generality of the approach for broad image sets.

**Figure 4.**
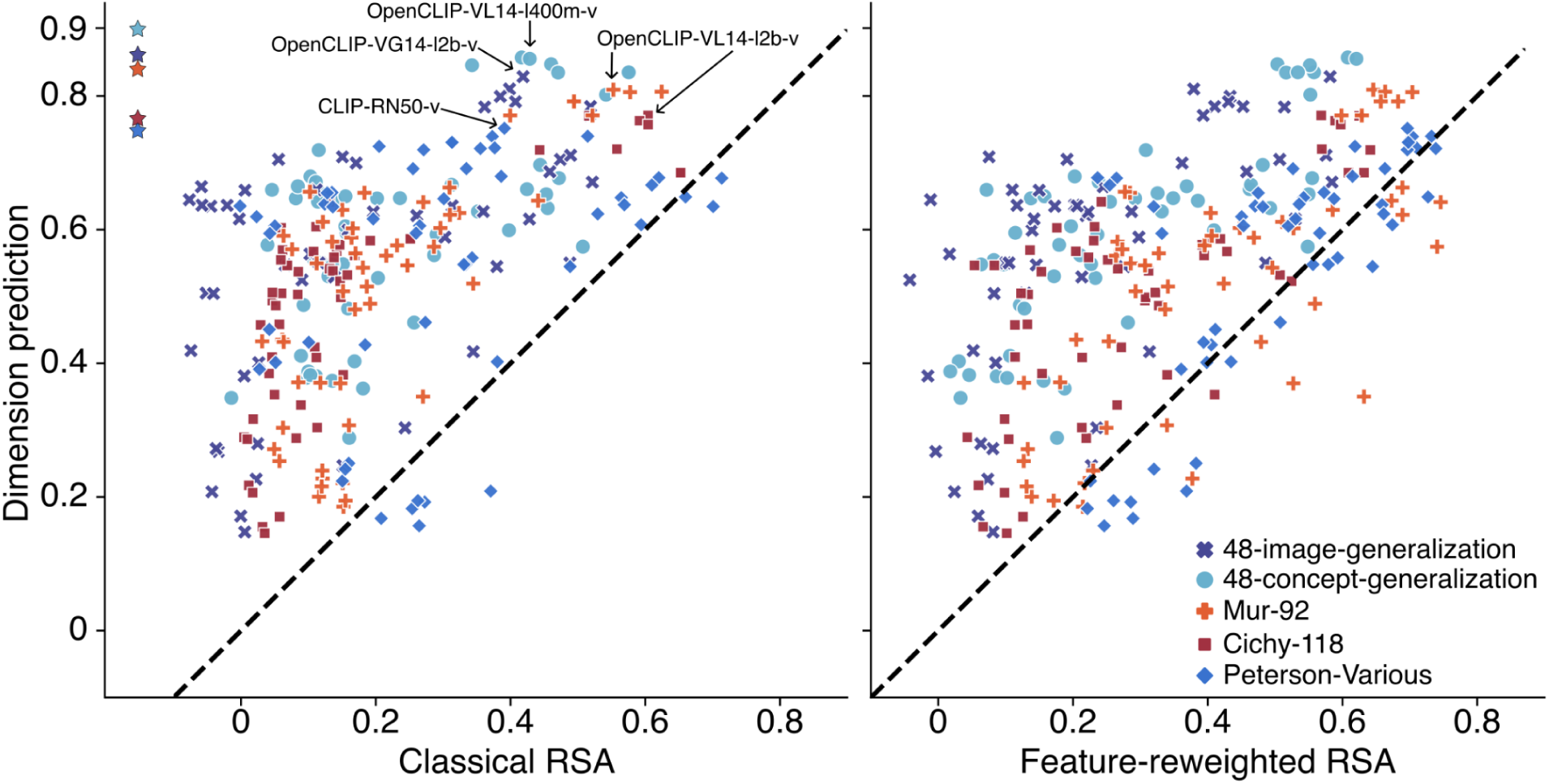
Predictive performance of all computational models for out-of-sample heterogeneous image sets. Our dimension prediction approach (DimPred) yielded similarity scores that correlated very highly with ground-truth similarity scores across five different heterogeneous validation sets. Across image sets, DimPred outperformed either classical (cRSA; left) or feature-reweighted RSA (FR-RSA; right) in 256 and 231 of 265 cases, respectively. The best computational model for each image set is indicated using the model abbreviation from Supplement 4. An ensemble DimPred model consisting of the best 8 computational models performed even better (denoted by a star symbol for each image set in the left panel).

### Humans can successfully rate images on embedding dimensions - but rely on information similar to those of DNNs

While DNN-based predictions yielded excellent performance, we asked whether humans could solve the prediction of dimensions with similar or better efficiency, and if so, whether combining humans with DNNs would lead to further improvements in predictive performance. To this end, we utilized crowdsourcing to gather human ratings of images on the 49 dimensions (see Figure 2, top). In the crowdsourcing task, after training, participants had to rate the images on each of the 49 SpoSE dimensions separately using a visual Likert-type scale. Based on these ratings, we computed an RSM which we then compared to a behavioral ground-truth RSM (for further details see *Methods*). As target images, we used the 48-image-generalization and 48-concept-generalization image sets.

Our results showed that humans were successful at providing sensible dimensional ratings of images; the resulting similarity scores correlated highly with ground-truth similarity scores for the two image sets (image set 1: *r* = 0.804, image set 2: *r* = 0.838) (see Figure 5 and Supplement 3). Next, we combined the human-based similarity scores with the similarity scores yielded by DimPred using a single neural network (OpenCLIP-RN50×64, OpenAI, visual) by averaging the predicted similarity scores of both. Compared to human-only predictions, this combination improved performance notably (image set 1: *r* = 0.861, *p*_diff_ = 0.002; image set 2: *r* = 0.884, *p*_diff_ = 0.002). Also in comparison to DNN-only predictions, performance was improved (original *r*s = 0.811 and 0.848, *p*_diff_ = 0.014 and 0.005, respectively). When combining the human-based similarity scores with the similarity scores yielded by the top-8 ensemble DimPred model, performance improved even more compared to human-only predictions (*r*s = 0.878 and 0.900, and *p*_diff_ < 0.001, respectively), while performance improvement was smaller and non-significant in comparison to ensemble-only predictions (image set 1: *r* = 0.861, *p*_diff_ = 0.869; image set 2: *r* = 0.898, *p*_diff_ = 0.244; all *p*-values uncorrected). This demonstrates that in comparison to the ensemble approach, combining DimPred with human predictions did not yield a statistically significant benefit, and the small improvement in absolute prediction accuracy may not justify the additional overhead of collecting human dimension ratings.

**Figure 5.**
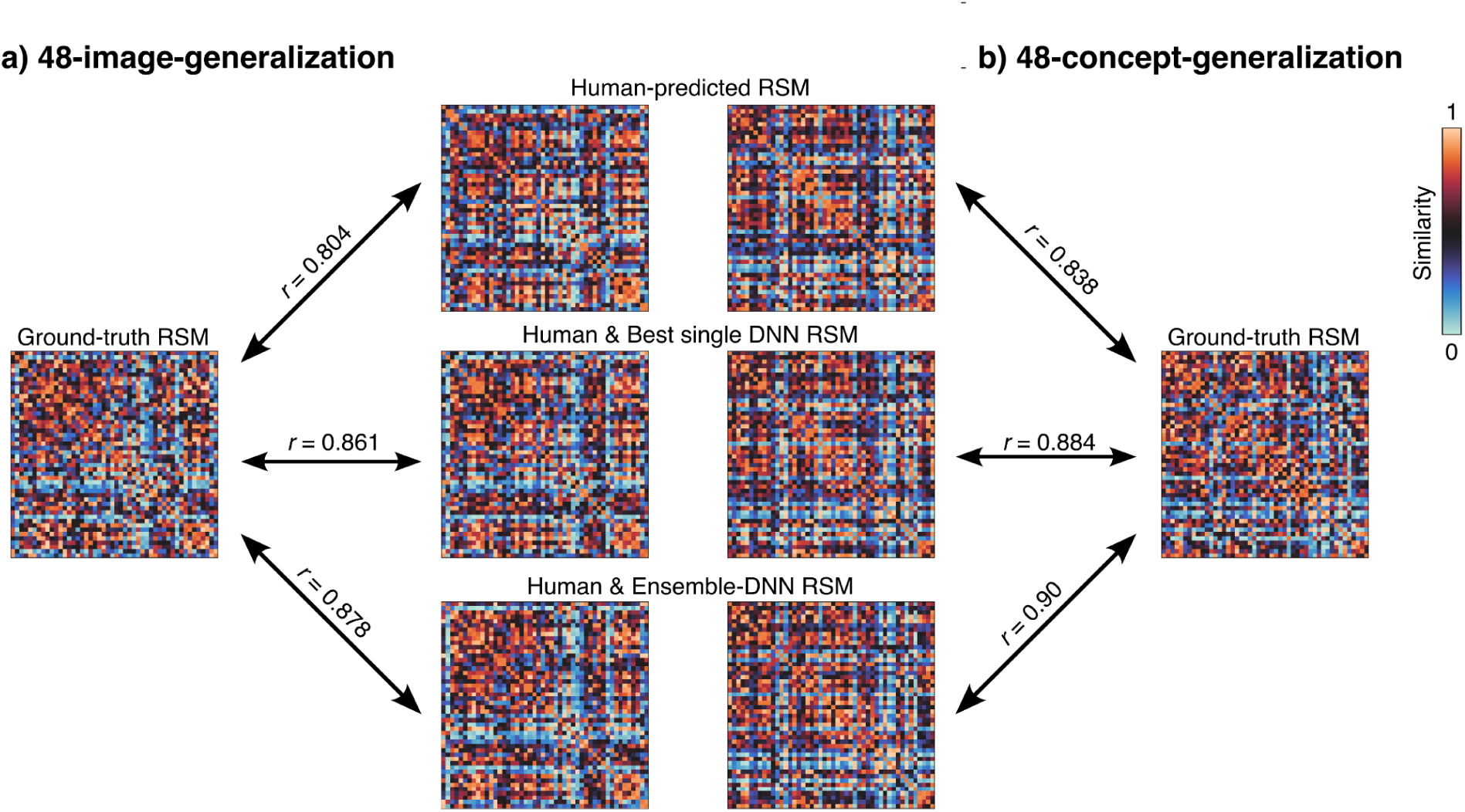
Correspondence between human-predicted, human & best single DNN combined or human & ensemble-DNN combined similarity matrices and ground-truth similarity matrices. For the two image sets a) the 48-image-generalization and b) the 48-concept-generalization image set, the representational similarity matrix (RSM) solely based on human ratings of image dimension values (top row), an average of that RSM and the best single DNN-predicted RSM (middle row), and an average of the human-based RSM and the top-8 ensemble DimPred model RSM (bottom row) was correlated with the respective ground-truth RSM. While humans provided sensible dimensional ratings of images evidenced by RSMs correlating highly with ground-truth RSMs, enriching them with RSMs yielded by DimPred for OpenCLIP-RN50×64 (OpenAI, visual) or with RSMs yielded by the ensemble DimPred model increased correspondence with ground-truth RSMs notably compared to only using human-derived RSMs.

### Predictive accuracy depends on granularity of an image set’s similarity structure

The image sets investigated so far contained many different object categories (e.g., aardvark, ambulance, bed). However, DimPred had been trained on similarity data sampled for a broad set of categories but with only very sparse sampling within higher-level categories (e.g., animals). We thus wondered what the limits of the performance of DimPred are when applying it to image sets from much more homogeneous image classes (e.g., pictures of furniture only). To test this, we utilized five more validation sets containing images from Peterson and colleagues^41^ that each consisted of 120 images of either exclusively animals, automobiles, fruits, furniture, vegetables and again used cRSA and FR-RSA as baseline approaches.

The respective results are presented in Figure 6. As expected, DimPred performed poorly on these image sets, with the ensemble model showing only slightly increased performance for most image sets. For all homogeneous image sets, DimPred was better only in 73 (Figure 6, left panel) or 59 (Figure 6, right panel) of 265 cases when compared to cRSA or FR-RSA as a baseline, respectively, reaching predictive accuracy of only up to *r* = 0.541.

**Figure 6.**
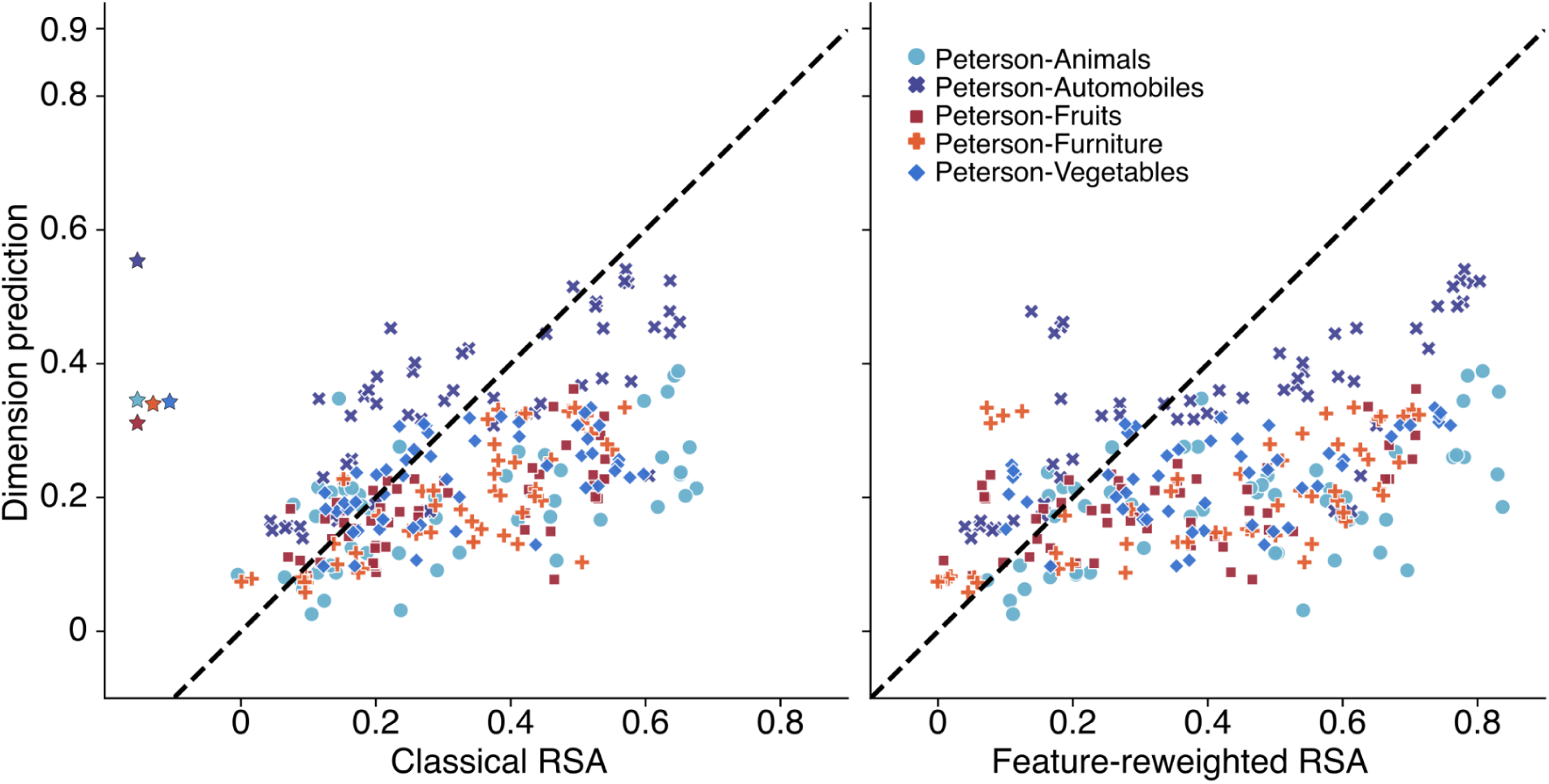
Predictive performance of all computational models for out-of-sample homogeneous image sets. Dimpred yielded similarity scores that did not correlate well with ground-truth similarity scores for five validation sets containing homogeneous images. Across image sets, DimPred was outperformed by either classical (cRSA; left) or feature-reweighted RSA (FR-RSA; right) in 73 and 59 of 265 cases, respectively. An ensemble DimPred model consisting of the best 8 computational models sometimes performed slightly better (denoted by a star symbol for each image set in the left panel).

### Revealing SPoSE-dimension specific areas in images important for similarity derivation

Despite the challenges of applying DimPred to the prediction of similarity judgments at a more fine-grained level, the prediction of similarity at scale for broadly-sampled natural images opens up many potential avenues. To showcase this potential, we first asked whether we could use the dimension predictions to highlight the likely image features underlying similarity judgments in humans. DimPred yields end-to-end prediction from images to individual dimensions, allowing us to observe the change in dimension prediction when occluding parts of the original image^58^. To this end, we used the iterative masking approach RISE^59^ that repeatedly occludes random image regions while observing the effect on the prediction performance of the model (see *Methods*). This approach allowed us to generate heatmaps that indicate the importance of image regions for a given SPoSE dimension. Beyond the prediction of individual dimensions, this approach allows us to compute an overall relevance score for the image regions involved in similarity judgments. Specifically, multiplying the predicted heatmaps with the dimension scores and averaging across dimensions provides us with a single heatmap of image region importance for similarity judgments. Example heatmaps are shown in Figure 7. For each of the example images, heatmaps for the global relevance as well as for the three most important image-specific SPoSE dimensions are shown. Indeed, the visualization method produces interpretable results, highlighting not only the image regions in line with the individual predicted dimensions but also which image regions are behaviorally-relevant for similarity and categorization judgments. Together, combining DimPred with visualization techniques offers a window into understanding which parts of an image may underlie its mental representations.

**Figure 7.**
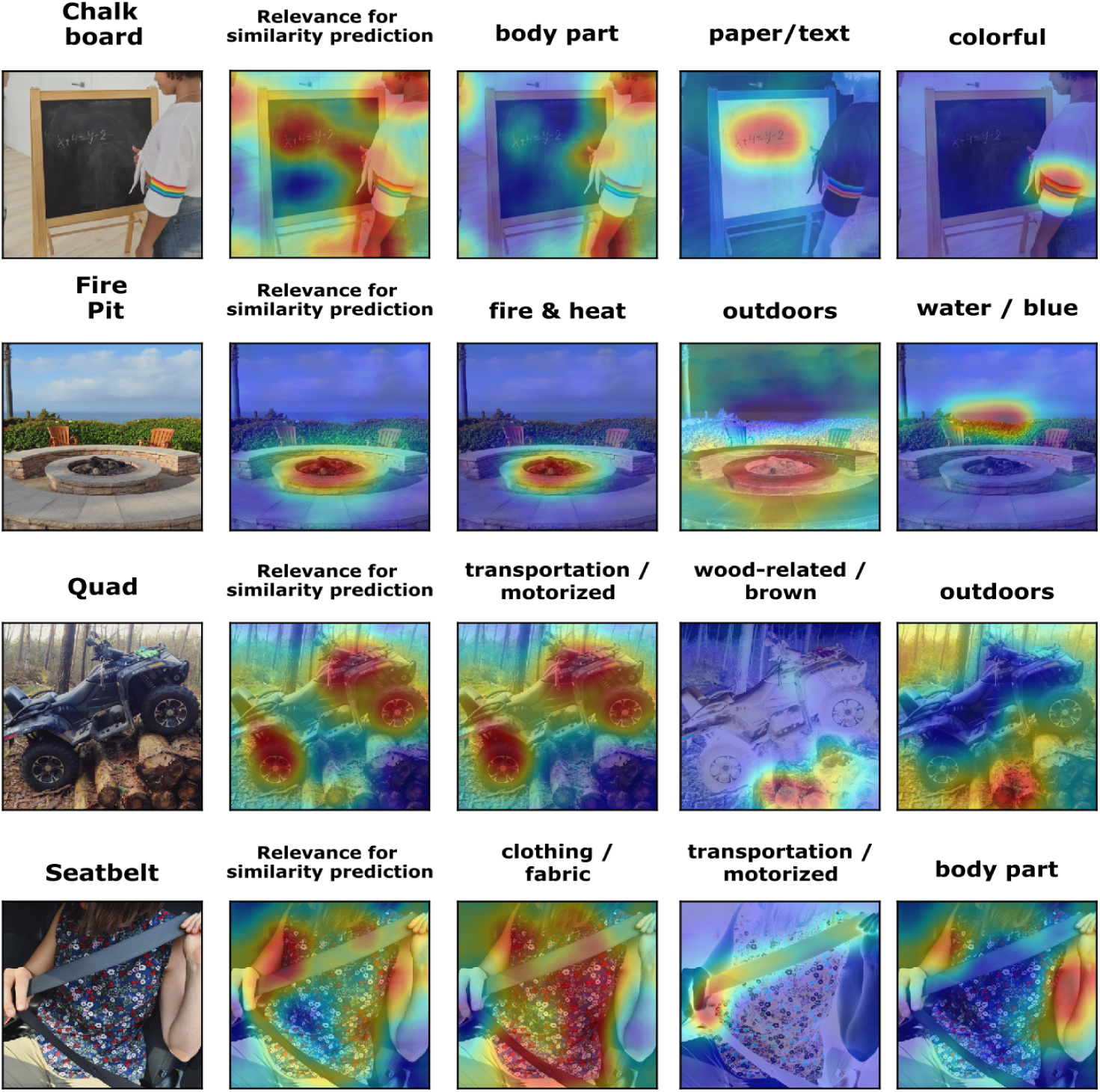
Visualization of image regions relevant for predicting different dimensions underlying similarity judgments. For each of the example images, the three most important SPoSE dimensions are shown. In addition, the relevance for similarity prediction reflects the weighted integration across all dimensions, highlighting candidate image regions that are behaviorally relevant for similarity judgments and categorization behavior.

### DimPred improves brain-behavior correspondence

To showcase the potential of DimPred for the application to external datasets, we applied DimPred to a publicly available functional MRI dataset of three participants who each saw 8,740 object images from 720 categories^60^. As a baseline, we fit a voxel encoding model of all 49 dimensions. Since dimension scores were available only for one image per category^36^, for the baseline model, we used the same value for each image of the same category and estimated predictive performance using cross-validation. To test the degree to which image-specific effects contributed to the prediction, we used DimPred to attain predicted dimension scores for each image and fit another encoding model that was evaluated separately. As a computational model, we used OpenCLIP-RN50×64. The relative improvement for each participant is shown in Figure 8. As can be seen, regions across the visual system show consistent improvements in predictive performance, demonstrating clear image-specific effects across the entire visual system, including high-level visual cortex. We have applied this approach in practice for the prediction of functional MRI^61^ and MEG data^62^. Together, these results highlight the potential of applying DimPred to existing dataset to gain novel insights into the functional organization of the visual system.

**Figure 8.**
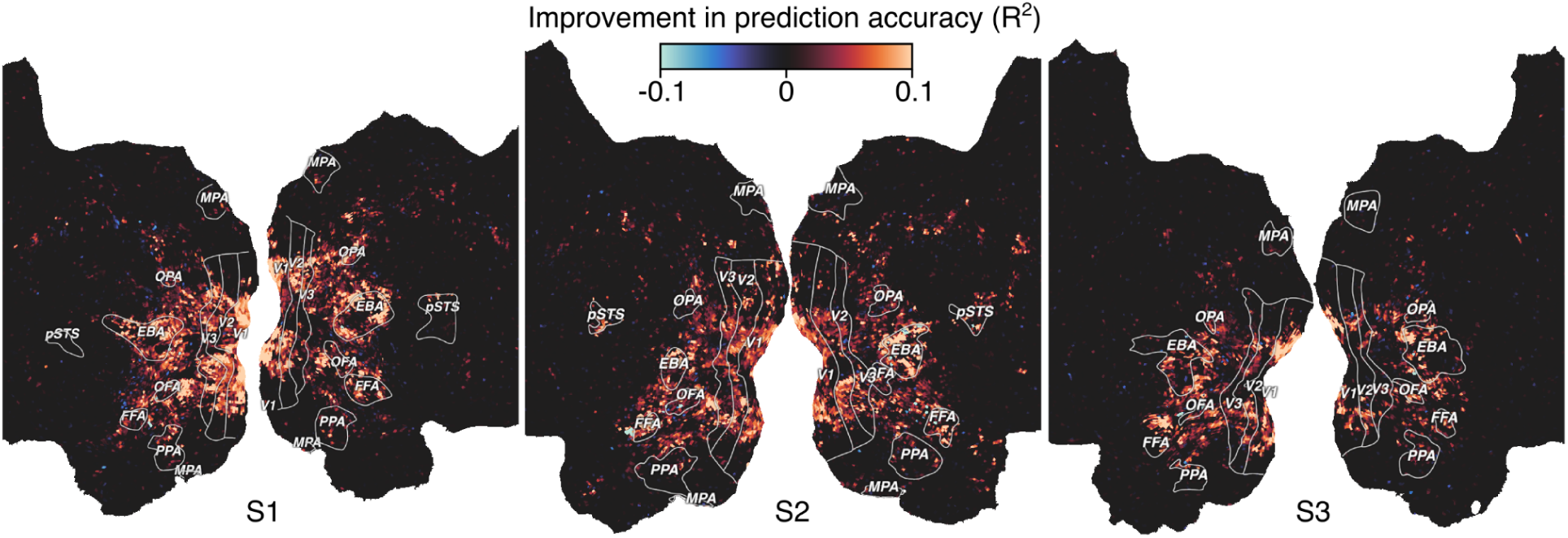
Prediction accuracy improvement of an fMRI voxel-wise encoding model based on the DimPred embedding. For three subjects, we computed an encoding model to predict voxel-wise fMRI activity, either using a coarser category-insensitive object image embedding or using the category-sensitive image-specific embedding as produced by DimPred based on activations from OpenCLIP-RN50×64. The relative improvement in encoding-model prediction accuracy when using DimPred is color-coded separately for each subject on a cortical flat map.

## DISCUSSION

Perceived similarity is central to understanding mental representations and how they relate to brain activity and behavior^1,2,6,9,13^. In addition, it plays a crucial role in evaluating computational models for their potential to explain neural or behavioral data^41^. In this study, we present an image-computable model of perceived similarity that uses the activations of a deep neural network (DNN) to predict meaningful similarity scores for thousands of object images. Our approach, DimPred, demonstrated a notable correspondence between the predicted similarity scores and ground-truth similarity across various image sets containing diverse categories. Given the high out-of-sample accuracy, this method suggests that predicted similarity scores can serve as a substitute for perceived similarity judgments of broadly-sampled, diverse object image sets, especially when the collection of such judgments is too effortful or costly. Our results highlight examples of how DimPred can be used in practice to yield new insights into the mental and neural representations of objects.

Much previous work combining artificial neural networks with human similarity judgments has focused on revealing the correspondence between artificial neural networks and human similarity judgments. Thus, explicitly or implicitly, many of these studies have attempted to automate the estimation of perceived similarity of stimuli. Most prior research fitted embeddings of off-the-shelf DNN layers directly to predict pairwise similarity judgments^35,37,38,41^, which required fitting all relevant features used in similarity comparisons simultaneously. In contrast, our dimension-based prediction framework DimPred breaks down this problem by separately predicting individual interpretable dimensions. In addition to the complexity of the prediction, much previous work relied on small similarity datasets^35,38,39,42,43^, raising questions about their generalizability for predicting perceived similarity. Our results demonstrate that DimPred’s predictive performance surpasses direct prediction approaches for datasets including broadly-sampled object categories and even generalized to datasets using other similarity tasks. In this context, it is important to note that many previous approaches, despite their seemingly impressive performance on existing datasets^35,38,41,42^, were not genuinely validated out-of-sample. Validation data often included several examples from the same object used during training, or performance was estimated in a non-independent manner by not leaving out objects but similarity judgments. In contrast, DimPred was evaluated both across category and across similarity tasks and continued to yield notable out-of-sample performance.

While DimPred showed strong performance on broad image sets, for within-category similarity prediction, performance was more limited. While this result might be due to the lack of relevant information in the neural networks used for training the approach, recent work successfully predicted representational dimensions for the category of manipulable objects, demonstrating that within-category dimension prediction from DNN activations is feasible^63^. This suggests that the performance of DimPred critically depends on the granularity of the similarity judgments from which the dimensions are derived. Since DimPred was trained on broad comparisons of many objects spanning diverse categories, it emphasized comparisons at a less fine-grained level than within-category comparisons. The recent success in predicting representational dimensions within categories^63^ highlights DimPred’s potential for predicting similarity ratings at varying levels of granularity. Future work could involve training representational embeddings at multiple levels, both within and across categories, to determine the ideal level of granularity for a given set of similarity comparisons, thereby maximizing predictive accuracy.

Predicting dimensions from human ratings yielded an accuracy very similar to that of DimPred. Combining both human and DNN ratings did not yield notable improvements over the performance found in an ensemble model of DNNs alone, but the addition of DNNs still slightly improved human ratings. This comparison might underestimate human performance, as human dimension ratings were based on a limited number of participants, which is typical of empirical studies. Nevertheless, the strong similarity in predictive accuracy between humans and DNNs demonstrates that DNNs captured similar information to that used by humans when rating dimensions. This indicates that much of the information needed to explain perceived similarity at the level of object images in this study may already be present in the best-performing DNNs.

The dimensions that span the representational space of objects capture both visual and semantic aspects of our visual world^36^. Interestingly, the best performing computational models were those that combined visual and semantic information (i.e., the OpenCLIP models) compared to classical deep neural networks that encode visual information only. Determining whether this performance is due to dataset size, dataset diversity, multimodal training, or the effective number of prediction targets is an important question for future work.^49,50,63^

Given the fact that DimPred can predict similarity judgments for broadly-sampled object image sets at scale, this opens the door to many novel applications, some of which we highlighted in the present work. First, in the past, it has remained unclear which image regions are used for similarity judgments, and only recent efforts have highlighted behaviorally relevant regions in images^64,65^. In the present work, we not only identified which image regions were potentially behaviorally relevant for similarity judgments. We also revealed what image regions were potentially relevant for individual representational dimensions (e.g., an animal-related dimension). Second, we generated image-specific predictions to improve encoding models of the visual system, which we highlighted with functional MRI data^61^ but which has also been applied to improve the prediction of MEG data^62^. This highlights that DimPred could be applied not only to future dataset but could even be applied post-hoc to already existing data, thus enabling others to answer new questions with already existing data. Furthermore, DimPred may allow teasing apart image-specific from general category-specific visual representations and thus reveal the distribution of visual and semantic features relevant for the representation of individual objects.

## METHODS

### Image sets

We used 11 different image sets that consisted of images either of broadly-sampled objects from diverse high-level categories or of images within individual superordinate categories (e.g., vegetables). The first three image sets were taken from the THINGS dataset^44^, which consists of 1,854 object categories and 26,107 images of these objects. For the first image set, we used the reference image of each THINGS object category, resulting in 1,854 images. The second image set (48-image-generalization image set) consisted of 48 THINGS images. The objects had been used in a validation set in a previous study^36^ but here we used different object examples to test the generalization of DimPred to novel images. These objects had been chosen to be semantically diverse, and thus we anticipated that their prediction would be more challenging than a random selection of objects. The third image set consisted of object concepts that were not part of THINGS (48-concept-generalization image set) and were chosen to test the generalization to novel images and novel objects. The fourth image set^54–56^ consisted of 92 images of natural and artificial objects as well as human and non-human faces and bodies (Mur-92 image set). The fifth image set^15,57^ consisted of 118 natural images (Cichy-118 image set). Both image sets were chosen to test DimPred’s generalization to another similarity task (object arrangement). The last six image sets^41^ consisted of 120 images of either animals, automobiles, fruits, furniture, vegetables or a mix of these categories and were chosen to test the performance of DimPred within category or, for the last image set, the generalization to another similarity task (pairwise ratings).

### Layer activations from chosen deep neural networks

We extracted activations from various deep neural networks (DNNs) as computational models for all image sets. For a detailed overview of the DNNs and the specific modules we used, please see Supplement 4. Overall, we deployed various kinds of models: fully connected layers of convolutional networks, convolutional layers from convolutional networks, unsupervised networks, transformers, and CLIP models. Where applicable, we used versions of the DNNs that had been pre-trained on the 1,000 ImageNet object classes^66^. After extraction, we flattened activation vectors. To ease computational load, for each vector exceeding 4,096 units, we used sparse random projection to reduce the dimensionality of the layer to 4,096.

### The DimPred approach

DimPred works in two steps. In the first step, an L2-regularized multiple multi-output regression is fitted to a given DNN module’s image activations to predict a value for each image on each of the dimensions of a similarity embedding derived from human behavior. In the second step, the predicted embedding is used to build a predicted similarity matrix. For step 1, as a prediction target we used the Sparse Positive Similarity Embedding (SPoSE)^67^, which consists of 49 human-interpretable visual and semantic dimensions across images of 1,854 natural objects. SPoSE was derived from 1.46 million triplet odd-one-out similarity judgments and has been shown to capture human similarity judgments in this task close to the human noise ceiling^36^. For the prediction of this embedding, for the 1,854 reference image set and a given computational neural network model, we first extracted activations for the images. We then split these image activations into ten folds for an outer cross-validation. Hyperparameter tuning was carried out using three-fold inner cross-validation on the training set. We used fractional ridge regression^68^, with 70 different L2 hyperparameter candidates linearly spaced between 0.1 (high regularization) and 1 (no regularization). We opted for L2 regularization for increased stability and to avoid overfitting. Since SPoSE dimensions were non-negative, predicted dimension scores below zero were set to zero. This led to a predicted SPoSE embedding. In step 2, we computed a predicted representational similarity matrix by predicting the choice probabilities for each pair of stimuli in the context of all possible combinations of triplets. Finally, we used Pearson correlation to relate the predicted similarity matrix with a ground-truth similarity matrix. For the other image sets, we used the entire 1,854 reference image set for training, determined the hyperparameter with cross-validation, and predicted the similarity matrices for external datasets accordingly. Since the Mur-92 and Cichy-118 datasets were based on dissimilarities sampled using the multiple arrangement task, in this case dissimilarities were derived by computing Euclidean distances from the embeddings directly. To facilitate future use of DimPred, we provide a toolbox to run DimPred in Python (https://github.com/ViCCo-Group/dimpred).

### Ground-truth similarity matrices

The similarity matrix for the 1,854 reference image set was taken from Hebart and colleagues^36^. Please note that, for simplicity, we refer to the similarity matrix derived from this embedding as “ground-truth”, even though this is only a predicted similarity. Since the performance of DimPred may reflect overfitting to the prediction of SPoSE rather than perceived similarity predicted by SPoSE, we validated DimPred out-of-sample with the other datasets. The ground-truth similarity scores for the 48-image-generalization image set and the 48-concept-generalization image set were sampled in a separate online behavioral study (see below). Ground-truth behavioral (dis-)similarity scores for the other image sets were available online or provided by the authors of the respective original article.

### Comparison to classical and feature-reweighted RSA

To put the performance of DimPred into perspective, we compared it to classical and feature-reweighted RSA (cRSA and FR-RSA, respectively). Both are established methods used frequently to relate representational spaces to each other. Classical RSA is computed based on the raw activations of a given computational model for a given image set. Representational similarity matrices (RSMs) are compared by flattening the upper or lower triangular part of the similarity matrix and correlating it to the respective part of the ground-truth similarity matrix. As a comparison metric, we used Pearson correlation. Mur-92 and Cichy-118, we computed Euclidean distance scores based on the raw activations. For all of the other image sets, we computed Pearson similarity scores based on the raw activations.

FR-RSA is similar to classical RSA but reweights individual features using cross-validation, with the aim to maximize the linear correspondence with another representational space. By combining reweighted feature-specific similarities, FR-RSA achieves much greater power when relating representational spaces to each other^45^. For the image sets Mur-92 and Cichy-118, we computed squared Euclidean distances within the predicting features. For all of the other image sets, we computed dot-product similarities within the predicting features.

### Behavioral experiments

We conducted two online behavioral experiments with human research participants. In the first experiment, we sampled dimensional values for images while in the second experiment, we sampled odd-one-out data for images.

### Experiment 1

#### Participants

We recruited 1,097 participants via Prolific (www.prolific.com) residing in the United States. Participants who were overly fast or with overly repetitive responses or who did not complete the task were excluded, leaving 763 participants (566 female, 189 male, 8 other, mean age: 29.42, range: 18 - 74, SD = 10.08). Participants were compensated with an equivalent of 3 GBP per session and provided informed consent as well as agreed to the data storage policy before beginning with the experiment. The experiment was approved by the ethics commission of Leipzig University’s Faculty of Medicine (211/21-ek) and we have complied with all relevant ethical regulations.

#### Design and Stimuli

The experiment was run online via Pavlovia^69^ within each participant’s browser. In the first experiment, we let participants rate images of natural objects on each of the 49 SpoSE dimensions separately. As test images, we used the 48-image-generalization and the 48-concept-generalization image sets (and a third image set containing 200 images, see Supplement 3) making it a total of 296 images that were rated by participants. Participants were allowed to take part several times by rating different dimensions. Most participants (83.49%) rated one dimension only. Dimensions were presented visually using Likert-type scales, with example images arranged along the scale based on their scores for each dimension. Participants were required to rate different images based on the presented dimensions.

The rating scale consisted of seven points, where the lowest point (“not at all”), the second to lowest point (“very low”) and the highest point (“very high”) were labeled. Additionally, we used anchor images to visualize each point of the scale. Those anchor images were sampled from the reference image set but were not part of the rated images or the validation sets. Each dimension scale was labeled with the dimension labels from Hebart and colleagues^36^. Participants were able to click anywhere on the scale. Scales were percentile-based and were later converted back to continuous dimension scores.

Participants were trained on the task and received feedback. During the test phase, each participant rated all 296 test images on a given dimension. The final dataset after participant exclusion consisted of an average of 19.31 ratings per dimension (946 dimension ratings across the 49 dimensions).

### Experiment 2

#### Participants

498 online workers (mean age: 41.2, SD = 12.7, 275 female, 223 male) from the crowdsourcing platform Amazon Mechanical Turk took part in this experiment. Participants received 0.1 USD per completed session and provided informed consent as well as agreed to the data storage policy before beginning with the experiment. The experiment was approved by the ethics commission of Leipzig University’s Faculty of Medicine (211/21-ek) and we have complied with all relevant ethical regulations.

#### Design and Stimuli

The experiment was conducted on Amazon Mechanical Turk using a triplet odd-one-out task. In each trial, participants were presented with three images of natural objects and asked to select the image that was least similar to the other two. Each session consisted of 20 trials, and participants could participate in as many sessions as they wished. The images were taken from the 48-image-generalization and 48-concept-generalization sets, but images from the two sets were not mixed. We collected a total of 69,200 trials, which covered all possible triplet combinations of both image sets twice. Participants whose responses were too repetitive or too quick were excluded, leaving 63,440 trials. From these triplet ratings, we computed similarity scores, which served as ground-truth similarity matrices for downstream analyses.

### Heatmap generation

To identify which image features were most relevant for predicting SPoSE dimensions, we used the image explanation method Randomized Input Sampling For Explanation (RISE^59^). RISE is an image occlusion technique for generating heatmaps that highlight the relevance of different image parts to an embedding dimension. In brief, RISE applies a series of random masks to the original image. Each mask probabilistically determines whether each pixel is preserved, reduced in intensity, or set to zero. These masked images are then used to predict the SPoSE embeddings. The differences between the predicted values for the masked images and the original image are recorded. By averaging the results from many such masks, we obtain a relevance score indicating each pixel’s contribution to each embedding dimension.

### Statistical analyses

We calculated Pearson’s *r* between predicted and ground-truth similarity scores and, where applicable, tested Fisher’s z transformed differences between Pearson’s *r* values (two-sided).

### Software packages

For the DimPred implementation and all ensuing analyses, we used Python (Python Software Foundation. Python Language Reference, available at http://www.python.org) as well as the following Python main libraries: fracridge^68^, joblib (https://github.com/joblib/joblib), matplotlib^70^, numba^71^, numpy^72^, pandas^73^, seaborn^74^, scikit-image^75^, scikit-learn^76^, scipy^77^ and thingsvision^78^. We used the FR-RSA toolbox to conduct FR-RSA^45^.

## Data availability

All result files pertaining to this study will be made available upon publication via an OSF repository (https://osf.io/jtekq).

## Code availability

All analysis scripts pertaining to this study will be made available upon publication via an GitHub repository (https://github.com/ViCCo-Group/dimpred_paper). The toolbox to run DimPred itself in Python is available via GitHub (https://github.com/ViCCo-Group/dimpred).

## Acknowledgements

We thank Oliver Contier for his help with setting up the encoding models, Johannes Roth and Magdalena Gippert for helpful discussions, and Joshua Peterson and Ian Charest for sharing the similarity datasets used for validation. This work was supported by a research group grant by the Max Planck Society awarded to M.N.H., the ERC Starting Grant project COREDIM (ERC-StG-2021-101039712) awarded to M.N.H., and the Hessian Ministry of Higher Education, Science, Research and Art (LOEWE Start Professorship to M.N.H. and Excellence Program “The Adaptive Mind”). Open access funding provided by Max Planck Society. The funders had no role in study design, data collection and analysis, decision to publish or preparation of the manuscript.

## Credit authorship contribution statement

Philipp Kaniuth: Conceptualization, Methodology, Software, Validation, Formal analysis, Investigation, Data curation, Writing – original draft, Writing – review & editing, Visualization

Florian P. Mahner: Formal analysis, Visualization, Writing – review & editing

Jonas Perkuhn: Investigation, Writing – review & editing

Martin N. Hebart: Conceptualization, Methodology, Resources, Writing – review & editing, Supervision, Project administration, Funding acquisition

## Materials & Correspondence

Correspondence and material requests should be addressed to Philipp Kaniuth or Martin Hebart

