## supplementary_material for "A high-throughput approach for the efficient prediction of perceived similarity of natural objects"

### Supplement 1 - Predictive power of individual computational models for separate SPoSE dimensions

We evaluated whether our automatic dimension prediction approach (DimPred) can successfully predict images' values on the 49 interpretable embedding dimensions. Therefore, we evaluated our approach on the training set (in a way that avoids overfitting, for details, see *Methods*) because the 1,854 reference image set contains a ground-truth Sparse Positive Similarity Embedding (SPoSE). We evaluated the predicted embedding by correlating all images' predicted values on a specific dimension with the images' ground-truth values on the same dimension, separately for each computational model. The results are shown in Figure S1 which depicts  the correlation of DNN predicted dimensions with the true dimensions. First, we found that, across computational models, the predictive performance was better for early than for late SPoSE dimension, possibly due to increased sparsity of later dimensions[1]. Second, we found a strong variability between different computational models. Amongst all models we tested, the Open-CLIP networks performed exceptionally well with OpenCLIP-RN50x64 (OpenAI, visual) being the best computational model, achieving a predictive performance for individual dimensions of up to $r = 0.964$.

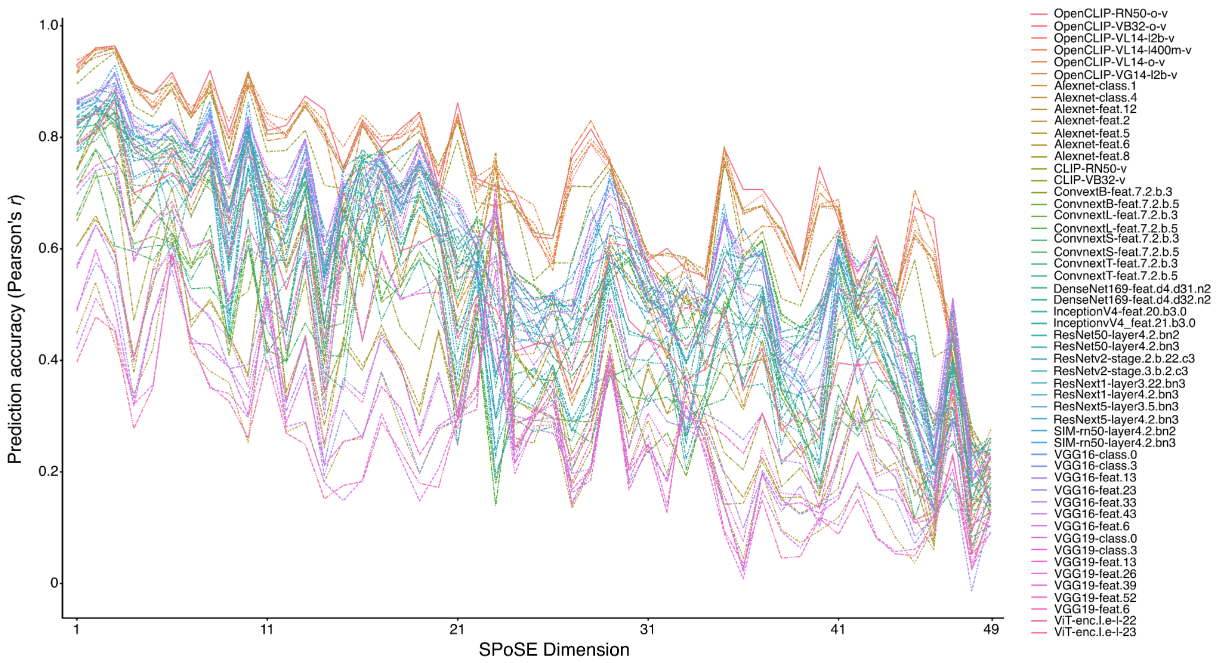


**Fig. S1. Predictive performance of all computational models for each predicted SPoSE dimension.** For the 1,854 reference image set, for each dimension (x-axis) separately, we correlated the predicted and ground-truth values (y-axis), separately for each computational model. For a mapping of computational model name abbreviations to full model names please see Table S4.

### Supplement 2 - Predictive accuracy of ensemble models

| | Best single model | 1 | 2 | 3 | 4 | 5 | 6 | 7 | **8** | 9 | 10 | 11 | 12 |
|---|---|---|---|---|---|---|---|---|---|---|---|---|---|
| 1854 reference | 0.888 | 0.895 | 0.914 | 0.922 | 0.923 | 0.925 | 0.925 | 0.925 | **0.926** | 0.926 | 0.925 | 0.922 | 0.92 |
| 48-image-generalization | 0.829 | 0.813 | 0.839 | 0.849 | 0.858 | 0.861 | 0.862 | 0.861 | **0.861** | 0.861 | 0.86 | 0.858 | 0.857 |
| 48-concept-generalization | 0.858 | 0.875 | 0.889 | 0.895 | 0.894 | 0.895 | 0.896 | 0.896 | **0.898** | 0.898 | 0.894 | 0.89 | 0.884 |
| Mur-92 | 0.809 | 0.772 | 0.824 | 0.834 | 0.832 | 0.83 | 0.837 | 0.841 | **0.843** | 0.843 | 0.841 | 0.84 | 0.837 |
| Cichy-118 | 0.771 | 0.774 | 0.772 | 0.78 | 0.778 | 0.782 | 0.779 | 0.77 | **0.768** | 0.765 | 0.761 | 0.757 | 0.751 |
| Peterson-Various | 0.752 | 0.687 | 0.722 | 0.73 | 0.733 | 0.737 | 0.739 | 0.744 | **0.751** | 0.759 | 0.762 | 0.76 | 0.76 |
| Peterson-Animals | 0.389 | 0.204 | 0.333 | 0.315 | 0.296 | 0.321 | 0.339 | 0.352 | **0.347** | 0.349 | 0.333 | 0.332 | 0.32 |
| Peterson-Automobiles | 0.541 | 0.54 | 0.551 | 0.535 | 0.536 | 0.542 | 0.543 | 0.548 | **0.55** | 0.551 | 0.548 | 0.544 | 0.54 |
| Peterson-Fruits | 0.362 | 0.287 | 0.291 | 0.284 | 0.294 | 0.297 | 0.298 | 0.305 | **0.312** | 0.313 | 0.304 | 0.304 | 0.299 |
| Peterson-Furniture | 0.335 | 0.294 | 0.311 | 0.318 | 0.331 | 0.326 | 0.342 | 0.337 | **0.341** | 0.347 | 0.339 | 0.345 | 0.341 |
| Peterson-Vegetables | 0.334 | 0.266 | 0.279 | 0.308 | 0.327 | 0.336 | 0.335 | 0.342 | **0.344** | 0.338 | 0.328 | 0.332 | 0.334 |

**Table S2. Ensemble model.** For the 1,854 reference image set, we combined the best *n* computational models (columns) separately for each predicted dimensional vector into an ensemble model. This ensemble model led to increased correlations between the resulting similarity scores it predicted and ground-truth similarity scores. The increase in this correlation continued up until combining around 8 different computational models per dimension. The success of this ensemble model found for the 1,854 reference image set carried over to the other out-of-sample image sets used in this study. For comparison, the first column denotes the performance of the best computational model as reported in the main text.

**Supplement 3 - Ability of humans to rate object images on separate SPoSE dimensions**

In addition to the 48-image-generalization and the 48-concept-generalization image set that were used in the first human experiments as described in the main text, we presented participants with an additional 200 images in the same experiment. We selected these 200 images from the 1,854 reference image set. The first 20 images were the same as those used to carry out object dimension ratings in a previous study[1]. For the remaining 180 objects, we chose a systematic, greedy selection strategy. Since our aim was to achieve a decent validation of the 49 SPoSE dimensions, we chose these images in a way that increased the variance across all 49 dimensions.

We assessed how well humans can rate images' values on the 49 interpretable embedding dimensions by correlating these 200 images' human-predicted values on each dimension with the images' ground-truth values on the same dimension. As can be seen in Figure S3, similar to one of the best computational models, the predictive performance was better for early than for late SPoSE dimensions. Overall, the participants' performance for individual dimensions reached up to *r* = 0.995 and was very similar to the best computational model's performance.

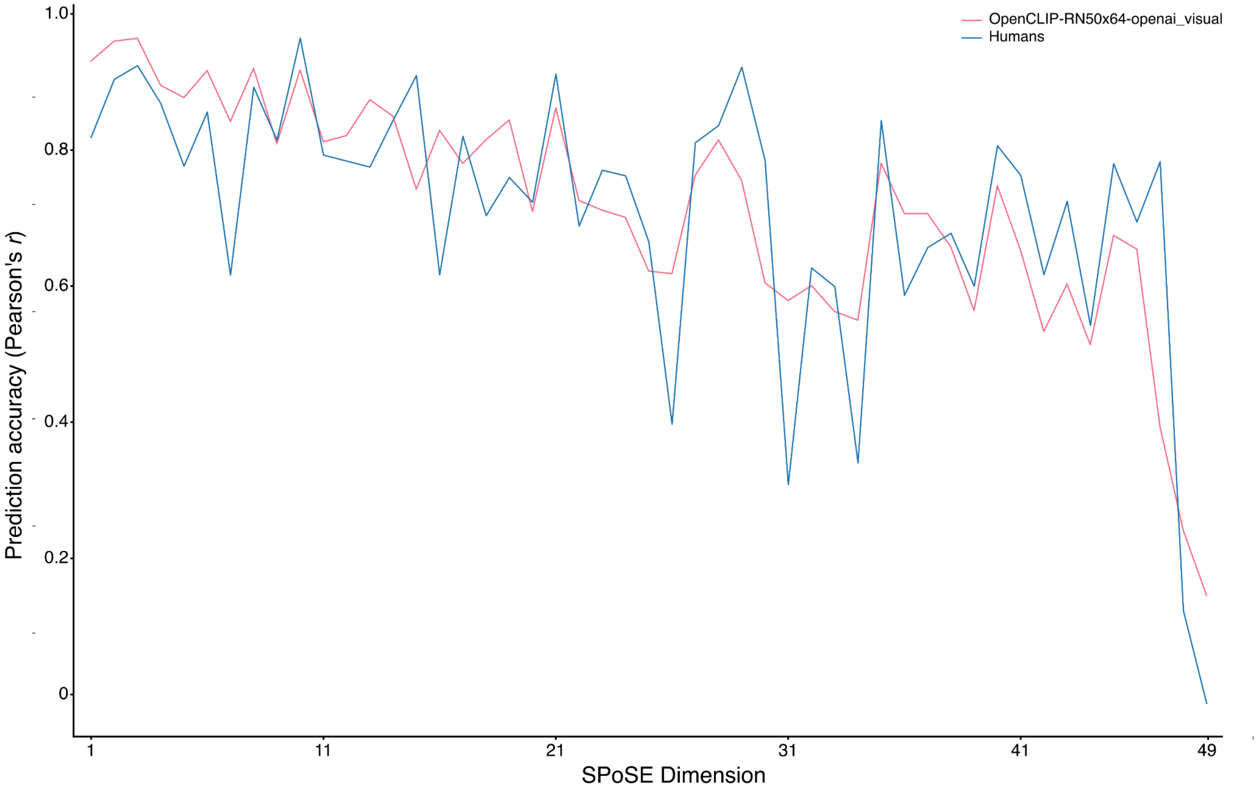


**Fig. S3. Association for each SPoSE dimension between ground-truth values and values as provided by human participants.** For 200 images (a subset of the 1,854 reference image set), we correlated the ground-truth values for each dimension (x-axis) separately with the respective values as provided by human participants. Similar to the best computational models, the association between ground-truth and human-provided dimensional values is better for early than for late SPoSE dimension.

### Supplement 4 - Overview of computational models used in this study

| Abbreviation example | Model (variant, dataset) | Modules | Reference |
|---|---|---|---|
| Alexnet-feat.2<br>Alexnet-class.1 | alexnet | features.2, features.5, features.6, features.8, features.12, classifier.1, classifier4 | 2 |
| CLIP-RN50-v | clip (RN50) | visual | 3,4 |
| CLIP-VB32-v | clip (ViT-B/32) | visual | 4,5 |
| ConvnextB-feat.7.2.b.3 | convnext_base | features.7.2.block.3,<br>features.7.2.block.5 | 6 |
| ConvnextL-feat.7.2.b.3 | convnext_large | features.7.2.block.3,<br>features.7.2.block.5 | 6 |
| ConvnextS-feat.7.2.b.3 | convnext_small | features.7.2.block.3,<br>features.7.2.block.5 | 6 |
| ConvnextT-feat.7.2.b.3 | convnext_tiny | features.7.2.block.3,<br>features.7.2.block.5 | 6 |
| DenseNet169-feat.d4.d31.n2 | densenet169 | features.denseblock4.denselayer31.norm2,<br>features.denseblock4.denselayer32.norm2 | 7 |
| InceptionV4-feat.20.b3.0 | inception_v4 | features.20.branch3.0,<br>features.21.branch3.0 | 8,9 |
| OpenCLIP-RN50-o-v | OpenCLIP<br>(RN50x64, openai) | visual | 4,10 |
| OpenCLIP-VB32-o-v | OpenCLIP<br>(ViT-B-32, openai) | visual | 4,10 |
| OpenCLIP-VL14-l2b-v | OpenCLIP<br>(ViT-L-14,<br>laion2b-s32b-b82k) | visual | 4,10,11 |
| OpenCLIP-VL14-l400m-v | OpenCLIP<br>(ViT-L-14,<br>laion400m-e32) | visual | 4,10,11 |
| OpenCLIP-VL14-o-v | OpenCLIP<br>(ViT-L-14, openai) | visual | 4 |
| OpenCLIP-VG14-l2b-v | OpenCLIP<br>(ViT-g-14,<br>laion2b-s12b-b42k) | visual | 4,10,11 |
| ResNet50-layer4.2.bn2 | resnet50 | layer4.2.bn2, layer4.2.bn3 | 3 |
| ResNetv2-stage.2.b.22.c3 | resnetv2_101x1_bitm | stages.2.blocks.22.conv3,<br>stages.3.blocks.2.conv3 | 3,12 |
| RexNext1-layer3.22.bn3 | resnext101_32x8d | layer3.22.bn3, layer4.2.bn3 | 13,14 |
| ResNext5-layer3.5.bn3 | resnext50_32x4d | layer3.5.bn3, layer4.2.bn3 | 14 |
| SIM-rn50-layer4.2.bn2 | simclr-rn50 | layer4.2.bn2, layer4.2.bn3 | 15 |

| | | | |
|---|---|---|---|
| VGG16-feat.6 | vgg16_bn | features.6, features.13, features.23, features.33, features.43, classifier.0, classifier.3 | 16 |
| VGG19-feat.6 | vgg19_bn | features.6, features.13, features.26, features.39, features.52, classifier.0, classifier.3 | 16 |
| ViT-enc.l.e-l-22 | vit_l_16 | encoder.layers.encoder-layer-22, encoder.layers.encoder-layer-23 | 5 |

**Table S4. List of all deep neural network models and their associated modules for which we extracted activations for our images.** The model and module names correspond to the parameters "model_name" and "module_name" in the thingsvision library[17] which was used to extract the activations. For some Clip and OpenClip models, the additional model parameter "variant" (and for OpenClip, additionally the "dataset") is denoted in parentheses. The first column gives examples for the abbreviations used in different figures.
